# Assessing mitochondrial function in angiosperms with highly divergent mitochondrial genomes

**DOI:** 10.1101/448902

**Authors:** Justin C. Havird, Gregory R. Noe, Luke Link, Amber Torres, David C. Logan, Daniel B. Sloan, Adam J. Chicco

## Abstract

Angiosperm mitochondrial (mt) genes are generally slow-evolving, but multiple lineages have undergone dramatic accelerations in rates of nucleotide substitution and extreme changes in mt genome structure. While molecular evolution in these lineages has been investigated, very little is known about their mt function. Here, we develop a new protocol to characterize respiration in isolated plant mitochondria and apply it to species of *Silene* with mt genomes that are rapidly evolving, highly fragmented, and exceptionally large (∼11 Mbp). This protocol, complemented with traditional measures of plant fitness, cytochrome c oxidase activity assays, and fluorescence microscopy, was used to characterize inter-and intraspecific variation in mt function. Contributions of the individual “classic” OXPHOS complexes, the alternative oxidase, and external NADH dehydrogenases to overall mt respiratory flux were found to be similar to previously studied angiosperms with more typical mt genomes. Some differences in mt function could be explained by inter-and intraspecific variation, possibly due to local adaptation or environmental effects. Although this study suggests that these *Silene* species with peculiar mt genomes still show relatively normal mt function, future experiments utilizing the protocol developed here can explore such questions in a more detailed and comparative framework.

## Introduction

Nearly all eukaryotes rely on mitochondria to supply the energy necessary for cellular function. Oxidative phosphorylation (OXPHOS) is the primary mechanism whereby mitochondria convert nutrients into cellular energy in the form of ATP. Energy conversion is accomplished via passing electrons from oxidized metabolic substrates, donated by high-energy reducing equivalents such as NADH, through a series of multisubunit protein complexes comprising an electron transfer system (ETS; a.k.a., electron transport chain; Fig. 1A). The resulting electron transfers generate a proton motive force across the inner mt membrane that is harnessed by ATP synthase to phosphorylate ADP. In many eukaryotes, oxygen acts as a terminal electron acceptor at cytochrome *c* oxidase (COX; Complex IV, CIV), resulting in a “coupling” of oxygen consumption and ATP synthesis. Accordingly, oxygen consumption by isolated mitochondria in the presence of ADP is routinely used as a proxy for OXPHOS function in diverse taxa (Balaban, 1990; Wilson *et al*., 1988; Chung *et al*., 2018; Haider *et al*., 2018;). ETS complex-specific substrates and inhibitors are also used to assess how different sites of electron entry exert control over OXPHOS function (Gnaiger, 2005). For example, rotenone (ROT) specifically inhibits Complex I (CI), allowing its contribution to total mt respiration to be parsed from other ETS complexes.

**Fig. 1.**
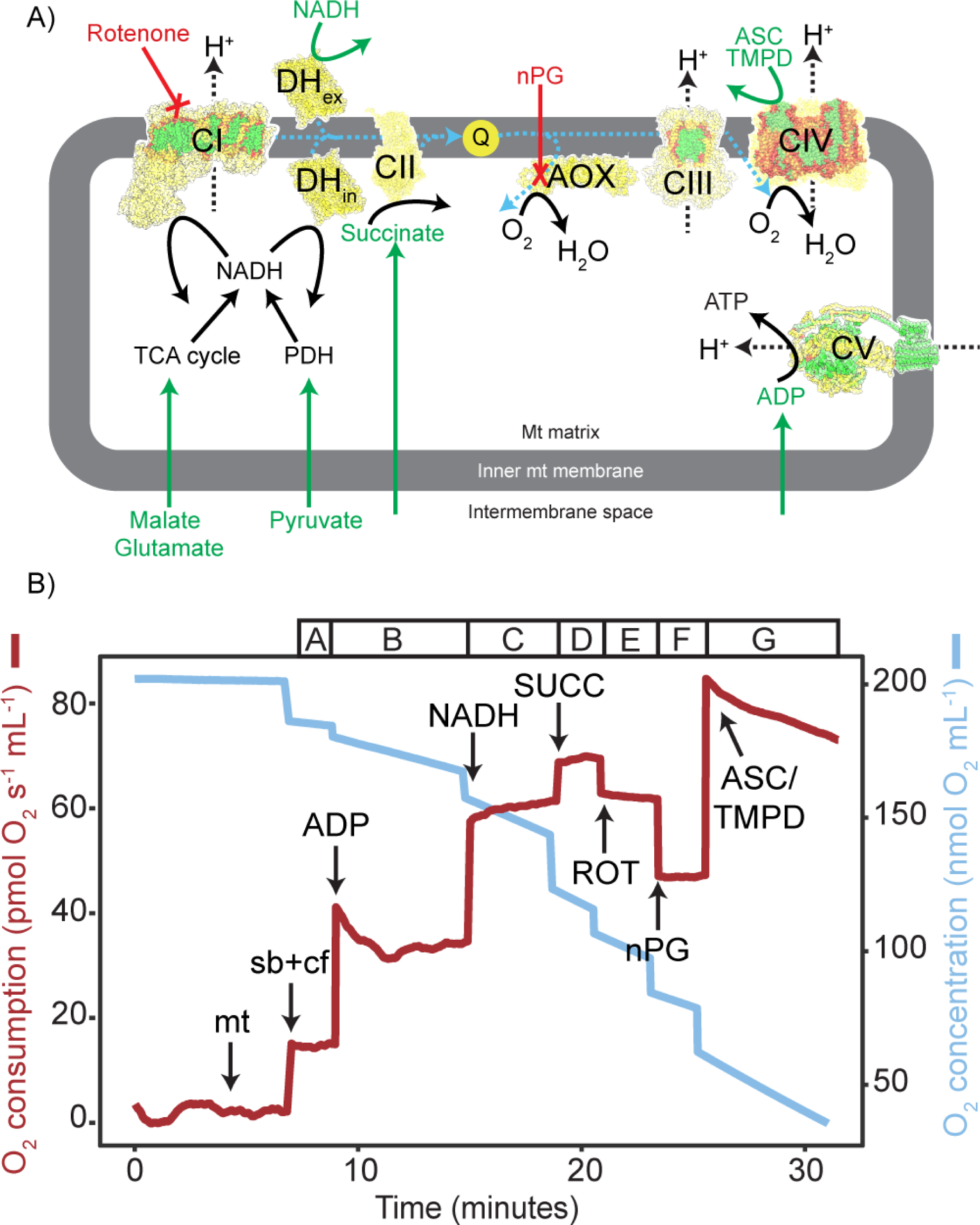
The plant mitochondrial electron transport system (A) and our protocol for quantifying mitochondrial respiration in seven different states (B). A) OXPHOS complexes are presented as structures from eukaryotic model species (PDB accessions 5LNK, 1ZOY, 1BGY, 1V54, 5ARA, 3VV9, and 4G6H). Residues are colored according to genomic identity in *S. conica*: nuclear-encoded residues are yellow, mt-encoded residues are green, and inter-genomic contact residues are in red (see Sharbrough *et al*., 2017 for details on identifying contact residues). Electron and proton flow are represented by blue and black dashed lines, respectively. Substrates added in our protocol are in green while inhibitors are red. B) Data were taken from a single sample that showed representative responses, although reoxygenation events and other details were removed to enhance clarity (a typical experiment lasted ∼2 hours). Black arrows indicate the addition of substrates/inhibitors: mt: mitochondrial isolate; sb+cof: the substrates and cofactors NAD+, TPP, CoA, malate, pyruvate, and glutamate; see main text for other abbreviations and details. Letters above the graph correspond to steps A-G in Table 1.

Such mt respirometry methods have been utilized in plants for decades (Ikuma and Bonner, 1967; Douce *et al*., 1977; Day *et al*., 1985). Many aspects of mt respiration in plants differ from mammals, making angiosperms useful study systems to understand OXPHOS (Affourtit *et al*., 2001; Millar *et al*., 2011; Jacoby *et al*., 2012). For example, although the mt ETS is more simplified than its bacterial ancestors (Berry, 2003), plant mitochondria contains a terminal alternative oxidase (AOX) not present in mammals (Fig. 1A). The AOX competes with CIV for electrons but does not result in proton translocation (Siedow and Girvin, 1980; Finnegan *et al*., 2004). Rather, AOX plays a role in preventing formation of reactive oxygen species and limiting oxidative stress (Siedow and Girvin, 1980; Considine *et al*., 2002; Fiorani *et al*., 2005; Watanabe *et al*., 2008). Alternative NADH dehydrogenases (i.e., internal and external NADH dehydrogenases, DH_in_ and DH_ex_; Fig. 1A), the coordination and balance of mt OXPHOS with photosynthesis, and the process of “photorespiration” also make plants interesting models to investigate mt function. However, such studies have largely used classic plant models such as Arabidopsis.

Of particular interest is the recent description of independent angiosperm lineages displaying abnormal patterns of mt genome evolution and structure (Mower *et al*., 2007; Sanchez-Puerta *et al*., 2017). In most angiosperms, mt genomes evolve slowly compared to the nuclear genome (Wolfe *et al*., 1987), whereas mammals show the opposite pattern (Brown *et al*., 1979). However, several independent angiosperm lineages have undergone recent and rapid accelerations in mtDNA evolution, resulting in a more animal–like balance between mt and nuclear substitution rates (Cho *et al*., 2004; Parkinson *et al*., 2005; Mower *et al*., 2007; Sloan *et al*., 2012a). Moreover, while most angiosperms mt genomes can be mapped as a single “master circle” chromosome (as in most eukaryotes)(Mower *et al*., 2012; Gualberto and Newton, 2017), multi-chromosomal mt genomes have also been found in several independent lineages (Alverson *et al*., 2011; Sloan *et al*., 2012a; Rice *et al*., 2013; Shearman *et al*., 2016; Sanchez-Puerta *et al*., 2017).

Recent evidence suggests that unusual angiosperm mt genomes can be associated with altered mt function. For example, the parasitic plant *Viscum* exhibits highly accelerated evolutionary rates, altered structure, and gene loss in its mt genome (Skippington *et al*., 2015, 2017) and was recently shown to: 1) have lost CI, 2) rely heavily on AOX and alternative NADH dehydrogenases, 3) have reduced abundances of all OXPHOS complexes, and 4) have an altered ETS configuration (Maclean *et al*., 2018; Senkler *et al*., 2018).

Certain species in the genus *Silene* are exemplars of angiosperm lineages with odd mt genomes, displaying: 1) accelerated rates of mt genome evolution (Mower *et al*., 2007; Sloan *et al*., 2009; Sloan *et al*., 2012a; Sloan *et al*., 2012b), 2) several dozen circular mt chromosomes (Sloan *et al*., 2012a; Wu *et al*., 2015), and 3) large expansions in mt genome size (up to 11 Mb) (Sloan *et al*., 2012a). While mt molecular evolution in *Silene* has been investigated thoroughly (Stadler and Delph, 2002; Mower *et al*., 2007; Sloan *et al*., 2009; Touzet and Delph, 2009; Sloan *et al*., 2012a; Sloan *et al*., 2012b; Sloan *et al*., 2014; Havird *et al*., 2015; Havird *et al*., 2017), it is entirely unknown how the curious features of their mt genomes have affected mt function.

Here, we investigate mt function in *Silene* using a novel method to comprehensively evaluate mt respiration by the sequential titration of multiple OXPHOS substrates and inhibitors in a single sample, complemented with measurements of plant fitness, COX enzyme activity, and live imaging of mitochondria. Through these experiments we addressed the following questions: 1) Do species with unusual mt genomic features have unusual mt function? 2) Does mt respiration show intra- or interspecific variation in *Silene*? and 3) Do correlations exist among different respiratory fluxes (i.e., a negative CI:AOX correlation as in *Viscum*), or between mt respiration and metrics of plant fitness?

## Methods

### *Silene* acquisition and growth

Four accessions of *Silene conica* were used: ABR (Abruzzo, Italy), SEN (Senez, France), KEWI (Wrocław, Poland, accession #8589), and KEWJ (Hungary, accession #1568) (described in Rockenbach *et al*., 2016). The first two were provided by collectors, while the latter two and a single sample of *S. subconica* (Gradevo, Bulgaria, accession #31398) were obtained from the Kew Millennium Seed Bank. The *S. subconica* accession was originally cataloged as *S. conica* but was morphologically similar to *S. subconica*, and phylogenetic analyses using transcriptomic data placed it with high support as sister to *S. subconica* to the exclusion of 19 *S. conica* populations (Havird *et al*., 2017). Therefore, we classify it as *S. subconica*. These accessions were propagated through several generations by self-fertilization to obtain seeds used here.

In December 2016, seeds were germinated on soil (Fafard 2SV mix supplemented with vermiculite and perlite) in SC7 Cone-tainers (Stuewe and Sons) on a mist bench under supplemental lighting (16-hr/8-hr light/dark cycle) in the Colorado State University greenhouse. Either eight (SEN, KEWI, and KEWJ), 16 (*S. subconica*), or 24 (ABR) seeds were used per sample. Seeds were obtained from at least two different parents and were planted in a randomized layout to minimize spatial effects. Germination success was scored three and four weeks after planting, after which seedlings were transferred off the mist bench and began to be treated with supplemental watering and fertilizer. Fifteen weeks after germination, the plants were transferred to 1 gal pots in order to promote vegetative growth. Plants were then monitored every day until the first flower was observed (∼6 months after germination).

### Mitochondrial isolation

Within 7 days after the first flower was obsered (5.6 ± 1.6 days standard deviation), mt respiration was quantified. Plants were therefore of similar developmental ages, although time between germination and experimentation varied by ∼2 months among individuals. Amount of vegetative growth also varied among individuals. For plants with enough tissue, 1 g of rosette and cauline leave were removed, whereas all rosette and cauline leaves were used for plants with less vegetative growth (minimum 0.1 g).

Leaves were finely diced and then ground with a mortar and pestle on ice in 5 mL of ice-cold mt isolation buffer (300 mM sucrose, 30 mM KH2PO4, 2mM EDTA, 0.8% polyvinylpyrrolidone, 0.05% cysteine, 5 mM glycine, 0.3% BSA, pH 7.5, with 15 mM-mercaptoethanol added just prior to use)(Delage *et al*., 2003). This homogenate was strained through four layers of cheese cloth and one layer of Miracloth, aliquoted into 1.5 mL microcentrifuge tubes, and centrifuged at 2790 g for 5 minutes at 4 °C to pellet large cellular debris and remove intact chloroplasts. The supernatant was then centrifuged at 12200 g for 15 minutes to pellet mitochondria. The mt pellet from each tube was then resuspended gently using paintbrushes in 100 uL of MiR05 buffer (110 mM sucrose, 20 mM HEPES, 10 mM KH_2_PO_4_, 20 mM taurine, 20 mM lactobionic acid, 3 mM MgCl_2_ 6H_2_O, 0.6 mM EGTA, 1 g/L BSA)(Gnaiger *et al*., 2000) and combined into a single tube for each sample. This suspension was centrifuged again at 12200 g for 5 minutes, the supernatant was removed, and the final mt pellet was resuspended in 200 uL of MiR05. This resulted in only a crude mt isolate and the final suspension was green, indicating the presence of broken plastids/thylakoids. However, further purification via Percoll or sucrose gradients and ultracentrifugation (e.g., following Murcha and Whelan, 2015) did not produce respiring mitochondria. Furthermore, the chloroplast-enriched isolates obtained after the first 2790 g centrifugation showed minimal respiration compared. Therefore, we proceeded using the crude mt isolation protocol.

### Mitochondrial respiration protocol

Mt respiration was quantified using an Oxygraph 2k (O2K) high-resolution respirometer (Oroboros Instruments GmbH, Innsbruck, Austria) from freshly obtained mt isolates. Respiration rates were normalized to total protein content, which was measured using a Qubit Fluorometer (ThermoFisher). Two O2Ks were used during experiments, each with two sample chambers, so that four samples were run simultaneously. Therefore, in a given day four samples were run immediately after mt isolation, while four samples were kept on ice until they were run (∼4 hours after mt isolation). Prior to adding the mt isolate, the oxygen content of MiR05 respiration buffer in the 2 mL chamber was air-calibrated to approximately 160 μM while stirring at 750 rpm, calculated using a barometer and known oxygen solubility of MiR05 (0.92). Following each experiment, respiration chambers were rinsed in 100% ethanol six times, immersed in 100% ethanol for at least 45 minutes, washed six times with 70% ethanol, and washed six times with ddH_2_o before adding MiR05 for the next experiment. All data were collected at 25 °C with lights turned off in the respiration chambers to minimize effects of photorespiration.

A primary aim of this study was to develop a comprehensive protocol for assessing the individual and integrative aspects of OXPHOS function in plant mitochondria. To this end, we created a multi-substrate and inhibitor titration protocol for plant mt isolates that generates seven distinct respiratory states suitable for individual and relative analyses (Fig. 1B). A detailed description of the protocol and associated respiratory states is provided in Table 1. Briefly, a volume of mt isolate with the equivalent of 0.25 mg of total protein was added to each respiration chamber, followed by a combination of substrates and cofactors (Table S1) that support mt NADH production and generate a low-flux “LEAK” respiration state facilitated by proton leak across the inner membrane in the absence of ADP (similar to “State 2” respiration in Jacoby *et al*., 2015; Fig. 1B, Table 1, Step A). ADP was added next to initiate OXPHOS, resulting in an increase in respiration rate due to dissipation of the proton gradient through the ATP synthase that limited electron flow in the preceding state (analogous to “State 3” respiration in Jacoby *et al*., 2015; Fig. 1B, Table 1, Step B). NADH was then added to support electron delivery to DH_ex_ (Fig. 1B, Table 1, Step C), followed by succinate (SUCC) to supply electrons via succinate dehydrogenase (Complex II, CII), generating the maximal OXPHOS-linked respiration rate observed in the experiment (Fig. 1B, Table 1, Step D). Rotenone (ROT) was then added to inhibit CI (Fig. 1B, Table 1, Step E), after which n-propyl gallate (nPG) was added to inhibit AOX (Fig. 1B, Table 1, Step F). Finally, ascorbate (ASC) then tetramethylphenylenediamine (TMPD) were added to fully reduce cytochrome c (CytC) and realize the maximal oxygen consumption (reductase) potential of CIV (Fig. 1B, Table 1, Step G), which typically exceeds that supported by upstream ETS and oxidation pathways (Gnaiger *et al*., 1998; Rossignol *et al*., 2003).

**Table 1.** High resolution respirometry titration protocol and associated oxygen flux states generated in mitochondrial respiration experiments. Abbreviations not listed above: ADP, adenosine diphosphate; CIV, Complex IV or cytochrome c oxidase; CoA, coenzyme-A; OXPHOS, oxidative phosphorylation; nPG, n-propyl gallate, an inhibitor of the alternative oxidase (AOX); Sb+Cf, substrates and cofactors supporting mitochondrial NADH production; TMPD, tetramethylphenylenediamine; TPP, thiamine pyrophosphate.

| Step<br>(Fig. 1) | Titration (mM) | Abbr<br>ev. | Site(s) of<br>electron entry | Explanation |
| --- | --- | --- | --- | --- |
| A | Malate (0.5)<br>Pyruvate (10)<br>Glutamate (10)<br>CoA (0.012)<br>TPP (0.2)<br>NAD<sup>+</sup> (2) | Sb+C<br>f | CI + DH<sub>in</sub> | LEAK-associated (non-coupled) respiration supported by saturating concentrations of substrates and cofactors for electron supply to Complex I and internal NADH dehydrogenases in the absence of ADP. Electron acceptors: CIV and AOX. |
| B | ADP (3) | ADP | CI + DH<sub>in</sub> | OXPHOS-associated respiration supported by electrons from Complex I and internal NADH dehydrogenases. Electron acceptors: CIV and AOX. |
| C | NADH (1) | NAD<br>H | CI + DH<sub>in</sub> +<br>DH<sub>ex</sub> | OXPHOS-associated respiration supported by electrons from all NADH dehydrogenases. Electron acceptors: CIV and AOX. |
| D | Succinate (40) | SUC<br>C | CII + CI +<br>DH<sub>in</sub> + DH<sub>ex</sub> | Maximal OXPHOS-associated respiration supported by electrons from CII and all NADH dehydrogenases. Electron acceptors: CIV and AOX. |
| E | Rotenone<br>(0.01) | ROT | CII + DH<sub>in</sub> +<br>DH<sub>ex</sub> | OXPHOS-associated respiration supported by electrons from CII and rotenone-insensitive NADH dehydrogenases (excluding CI). Electron acceptors: CIV and AOX. |
| F | *n*-propyl gallate<br>(0.5) | nPG | CII + DH<sub>in</sub> +<br>DH<sub>ex</sub> | OXPHOS-associated respiration supported by electrons from CII and rotenone-insensitive NADH dehydrogenases. Electron acceptor: CIV |
| G | Ascorbate (10)<br>TMPD (0.3) | ASC/<br>TMP<br>D | CIV + CII +<br>DH<sub>in</sub> + DH<sub>ex</sub> | Maximal capacity of CIV-mediated respiration supported by artificial electron donors. Electron acceptor: CIV |

Oxygen concentration was recorded every two seconds and oxygen flux was calculated using the previous 10 seconds of data. Generally, oxygen flux stabilized during each respiration state for several minutes and flux data were extracted from this stable region. However, after addition of ADP (Fig. 1B), it was consistently observed that respiration rates peaked, decreased, and then slowly returned to peak levels after ∼30 minutes (e.g., Fig. S1). Therefore, peak respiration immediately after adding ADP was used. Moreover, after adding ASC/TMPD, respiration peaked and then consistently declined (Fig. 1B), so peak respiration immediately after adding ASC/TMPD was used. Given the extended time needed to complete the protocol (∼2 hours), during the couse of some experiments (29%), oxygen was sufficiently depleted (< 10 nmol O2 mL^-1^) that chambers had to be reoxygenated by exposing them to air before continuing. Reoxygenation was never performed more than twice during any experiment, and preservation of mt quality was confirmed by stable retention of the respiration rate observed prior to reoxgenation.

Substrates and inhibitors were generally stored at −20 °C when not in use, with the exception of pyruvate that was made fresh before each experiment.

### Respiratory flux control factors

A primary advantage of multi-substrate/inhibitor titration respirometry protocols is the ability to calculate internally-normalized flux control factors (FCFs) that express specific aspects of respiratory control by normalizing flux to a common reference state (Pesta and Gnaiger, 2012). Several factors were calculated to represent the extent of respiratory control exerted by specific sites of electron entry and removal from the ETS (Table 2). The respiratory control ratio (RCR) was also calculated as a common measure of mt quality (Jacoby *et al*., 2015) by dividing the rate of respiration after adding ADP (Fig. 1B, Table 1, Step B) by the rate of respiration prior to adding ADP (Fig. 1B, Table 1, Step A).

**Table 2.** Respiratory flux control factors *Calculations refer to recorded rates of O2 flux at corresponding steps A-F in Table 1 and Fig. 1.

| Flux control factor | Calculation* | Explanation |
| --- | --- | --- |
| OXPHOS coupling efficiency | $1-(A/B)$ | Extent of ADP control over respiration, relating to the coupling efficiency of oxidative phosphorylation ranging from zero (completely non-coupled or no ADP control) to 1.0 (maximally coupled or 100% ADP control). |
| $DH_{ex}$ flux control | $1-(B/C)$ | Proportion of total NADH-supported OXPHOS-linked respiration contributed by external NADH dehydrogenases. |
| CI flux control | $1-(E/D)$ | Proportion of maximal (NADH + Succinate-supported) OXPHOS-linked respiration contributed by CI. |
| CII flux control | $1-(C/D)$ | Proportion of maximal (NADH + Succinate-supported) OXPHOS-linked respiration contributed by CII. |
| AOX flux control | $1-(F/E)$ | Proportion of OXPHOS-linked respiration mediated by the alternative oxidase (AOX). |
| Excess capacity of CIV | $(G/F)-1$ | Apparent excess respiratory capacity of cytochrome c oxidase over OXPHOS-linked respiration supported by CII and rotenone-insensitive NADH dehydrogenases. |

### Investigating inhibitory malate concentrations

Most of the substrate concentrations used above are commonly used in plant mt respiration protocols (e.g., Jacoby *et al*., 2015). However, we investigated the effects of varying malate concentrations because the typical concentrations of 10-30 mM used in previous plant experiments (e.g., Douce *et al*., 1977; Siedow and Girvin, 1980; Day *et al*., 1985; Jacoby *et al*., 2015) have been shown to inhibit mt respiration in other systems (Sumbalova *et al*., 2014a; Makrecka-Kuka *et al*., 2015). Two preliminary experiments were performed to assess inhibitory effects of high malate concentrations.

In the first experiment, respiration was examined in isolated mitochondria from *S. noctiflora* (a close relative of *S. conica* that also has an unusual mt genome) and *Pisum sativum* (a model for plant mt respiration with a more “normal” mt genome). Germination, mt isolation, and mt respiration were performed essentially as described above, except that additional malate was titrated into the chamber to evaluate responses to cumulative concentrations of 0.5 mM (as used above), 1 mM, 1.5 mM, 2 mM, and 3 mM. In the second experiment, a similar strategy was followed, with three differences: ADP was added before the addition of any malate, the AOX was inhibited at the start of the experiment by nPG, and malate concentrations ranged between 0.5 mM and 25 mM. In both experiments, additional substrates and inhibitors were added after the highest concentration of malate was achieved.

### COX enzyme activity

To complement measurements of mt respiration, COX enzyme activity was quantified in the same mt isolates. The assay is based on the oxidation of reduced CytC by COX in the sample, with the formation of oxidized CytC resulting in a linear decrease in absorbance at 550 nM (Storrie and Madden, 1990). Reactions were performed in 96 well plates using 200 L of sample buffer (120 mM Tris-KCl and 250 mM Tris-sucrose). Isolates were stored at −20 °C and 1-10 L of thawed stock mt isolate was added to each reaction, resulting in adding 1-16 ug per reaction. The reaction was started by adding 10 L of reduced CytC (0.22 mM) to each well (prepared by adding 5 L of 100 mM DTT per mL of oxidized CytC, mixing gently, and incubating at room temperature for at least one hour). The plate was then loaded into a Spectramax 2e spectrofluorometer plate reader (Molecular Devices, San Jose, CA), shaken for 10 seconds, and absorbance at 550 nm was recorded every 30 seconds for 10 minutes. The absolute slope of a line fitted through these data was taken as COX enzyme activity, which was normalized to protein concentration and averaged among two technical replicates.

### Metrics of plant fitness

Within seven days after flowering (1.7 ± 2.3 days standard deviation) but before tissue was harvested to prepare mt isolates, three metrics of plant fitness were quantified. Rosette diameter was measured to the nearest centimeter by measuring combined length of the largest rosette leaves. Plant height was measured similarly by using the tallest inflorescence. The number of stems per plant was also quantified. Finally, number of capsules and an approximation of total seed weight were quantified. These last two measures were performed after plants had senesced (about a year after flowering) and therefore: 1) these measures were not able to be quantified for all individuals due to some individual plant loss, and 2) these metrics were measured well after tissue had been removed for preparing mt isolates. It is likely that removing tissue therefore affected seed production, especially for plants with low amounts of vegetative growth. However, plants with little vegetative growth usually only had 1 or 2 stems and few buds at time of first flowering, suggesting these individuals would have produced few seeds regardless.

### Imaging mitochondria *in vivo*

Mitochondria were visualized using real-time imaging in living tissue of three species: *S. noctiflora*, which has unusual mt genomic features similar to *S. conica*, and *S. latifolia* and *S. vulgaris*, both of which show mt genomes more typical of angiosperms. Intact 7 to 10 day-old seedlings were incubated in the red fluorescent potentiometric dye tetramethylrhodamine methyl ester (TMRM) that accumulates reversibly in mitochondria in response to the inner membrane potential (Brand and Nicholls, 2011). Following incubation in freshly prepared 50 nM TMRM for 15 minutes, seedlings were mounted in TMRM between slide and cover slip within a chamber created using thin double-sided adhesive tape (Ekanayake *et al*., 2015). Microscopy was performed using a Zeiss Axioimager Z2 equipped with a Plan-Apochromat 100x/1.40 oil immersion objective and 63HE filter set (Ex: 572/25 nm, Em: 629/62 nm). Excitation was provided by a Zeiss HXP-120 metal halide lamp. Images were captured using a Hamamatsu Orca Flash 4 CMOS camera connected to Zeiss Zen software. Independent observations of at least three seedlings were made on at least two different dates.

### Statistical analyses

All statistical analyses were performed using R v3.4.3 (R Development Core Team, 2012) using general linear models and the *lm* function for continuous data and generalized linear models and the *glm* function/Poisson regressions for count data. Tukey post-hoc analyses were used to test for significant differences among populations.

## Results

### Germination rate and plant fitness

Three weeks after planting, all eight seeds had germinated for *S. conica* KEWI and KEWJ, while 7/8 had germinated for SEN, 15/24 for ABR, and 10/16 for the single *S. subconica* accession (Fig. 2A). These counts remained unchanged by four weeks after planting, and dormant seeds were excluded from the rest of the study. All seeds that germinated eventually went on to flower except for a single *S. subconica* individual that had not produced a reproductive stem by seven months after germination and was used for mt respiration and COX activity experiments but not scored for plant fitness metrics.

**Fig. 2.**
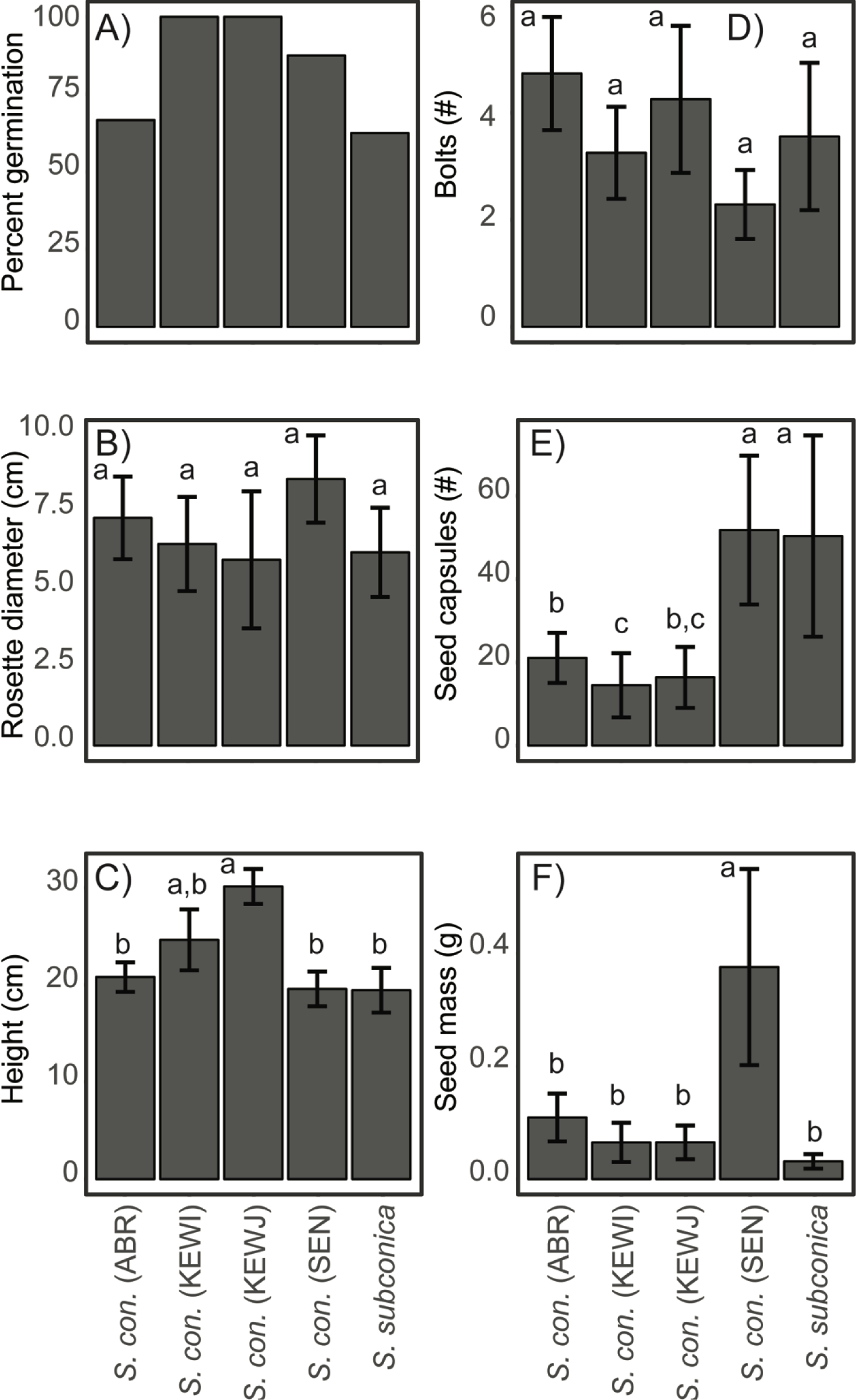
Fraction of seeds that germinated and metrics of plant fitness for the four accessions of *S. conica* and the single accession of *S. subconica* investigated: A) Percent seeds germinated, B) maximal rosette diameter, C) maximal height, D) number of stems, E) number of capsules, and F) total mass of seeds produced. Lowercase letters indicate significant groupings among accessions at *P* < 0.05 based on Tukey post-hoc tests. *n* = 7-15 for A) − D) and *n* = 5-13 for E) and F). Error bars show SEM.

There were few significant intra- or interspecific differences in common metrics of plant fitness. Although often correlated (Table S2), the differences in fitness metrics among accessions were not always congruent (Fig. 2B-F). For example, *S. conica* SEN had a total seed mass that was nearly four times larger than other accessions on average (*P* ≤ 0.044) but was also shorter and tended to have fewer stems (Fig. 2). Because many seeds were likely dropped naturally before seed mass was quantified, the number of capsules produced may be a more accurate metric of fitness, although the two metrics are highly correlated (Fig. S2). *S. conica* SEN as well as the single accession of *S. subconica* produced more than twice as many capsules as the other accessions (Fig. 2E, *P* < 0.001), but variation in these accessions was also much larger than in others, and significant differences may be driven by outliers with large numbers of capsules (e.g., in *S. subconica* zero to 128 capsules were prodcued).

### Developing a protocol to quantify mt respiration

During preliminary experiments to develop our protocol, we found that malate concentrations of 2 mM or higher in *S. noctiflora* and 1.5 mM or higher in *P. sativum* began to inhibit respiration (Fig. S3A, B). Adding succinate partially restored respiration in *P. sativum* but had little effect in *S. noctiflora* (Fig. S3A, B). In another experiment, malate concentrations above 3 mM decreased respiration, ultimately causing respiration to drop below baseline levels at 15 mM (fig. s3C, D). While adding NADH to supply electrons via DH_ex_ partially rescued respiration under these conditions (fig. s3C, D), adding succinate did not increase respiration in either species, consistent with an inhibitory effect of oxaloacetate production (from malate dehydrogenase) on CII (Makrecka-Kuka *et al*., 2015; Sumbalova *et al*., 2014b). Higher malate concentrations resulted in decreased respiration both when AOX was functional (fig. s3A, B) and inhibited (fig. s3C, D). Based on these results, we used 0.5 mM malate in our final protocol (Fig. 1B), and recommend that future studies carefully evaluate malate concentrations to optimize conditions for mt respirometry.

We ultimately developed a protocol that quantifies seven distinct mt respiratory states dependent on various components of the plant mt ETS (Fig. 1, Table 1). When this protocol was applied to isolated mitochondria from *Silene* species with unusual mt genomes, expected responses to respiratory substrates and inhibitors were observed (Fig. 3A). Although other species were not examined thoroughly, preliminary experiments using two closely related species with “normal” mt genomes (*S. latifolia* and *P. sativum*) yielded qualitatively similar results (fig. s3).

**Fig. 3.**
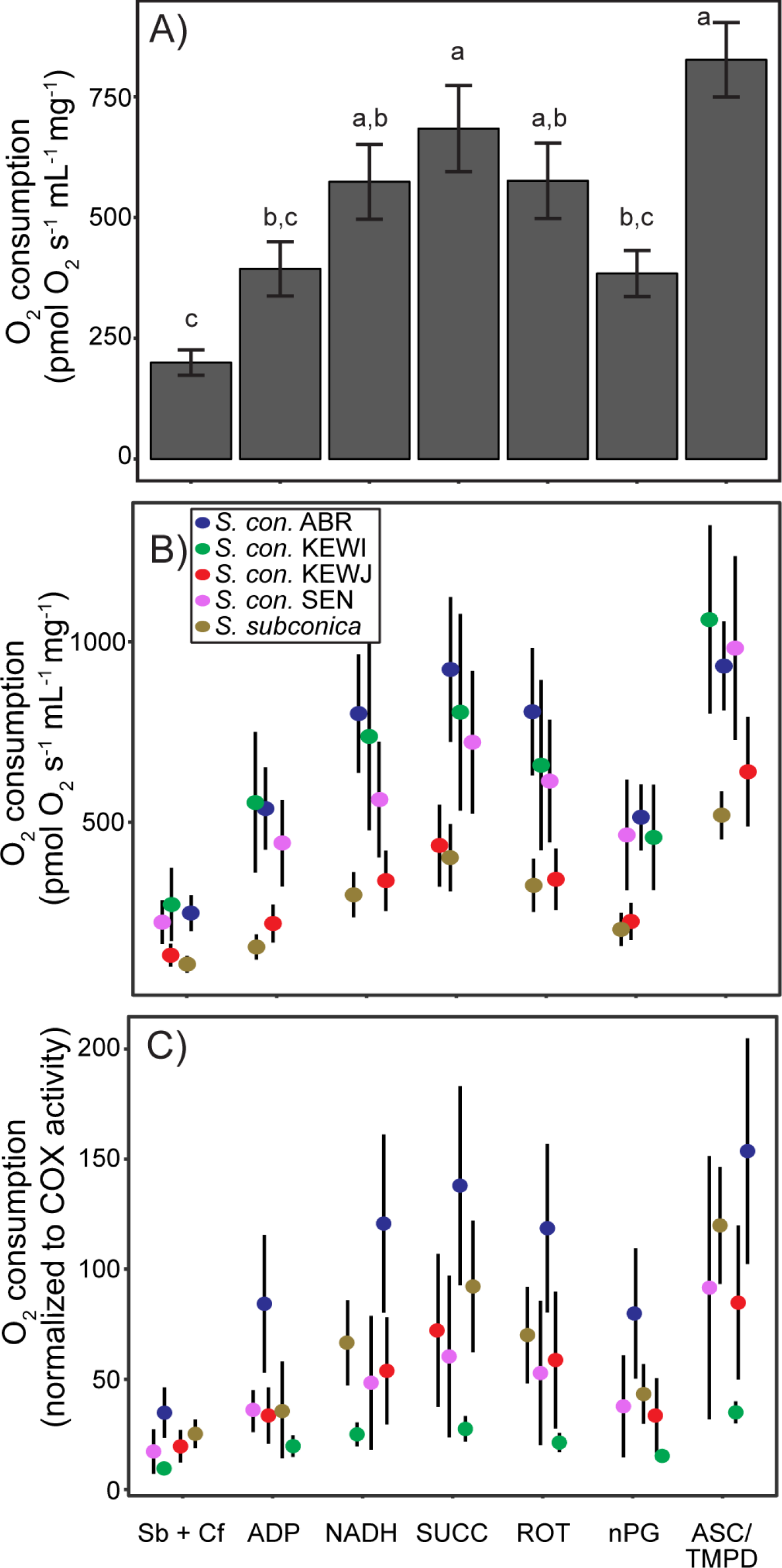
Respiration in isolated mitochondria of *Silene* during seven unique respiratory states. The seven respiratory states and their abbreviations follow steps A-G in Fig. 1B and Table 1. A) All accessions are pooled and respiration is normalized to mt protein input. B) Variation in protein-normalized respiration among accessions. C) Respiration normalized to COX activity. Lowercase letters indicate significant groupings among states at *P* < 0.05 based on Tukey post-hoc tests. There were no significant differences among accessions in B) or C). Error bars show SEM. *n* = 48 for A) and *n* = 7-15 for B) and C).

### Inter-and intraspecific variation in *Silene* mt respiration and COX activity

The RCR serves as a general metric for mt quality, with RCR values > 2.5 indicating high quality mt isolates (Jacoby *et al*., 2015). Our crude mt isolations from *Silene* had RCRs ranging from 1.1 to 3.9, with some of this variation explained by inter-and intraspecific differences (Fig. s4). For example, isolates from *S. subconica* and **S. conica** KEWJ had significantly lower RCRs than two of the **S. conica** accessions (Fig. s4). When samples that had RCRs less than 1.5 were excluded, the qualitative difference remained with *S. subconica*, but disappeared for KEWJ, mainly because all *S. subconica* samples had low RCRs (max = 1.9), while some KEWJ samples had higher RCRs (max = 2.5).

Intra- and interspecific differences in mt respiration were not significant during any of the seven respiratory states when respiration was normalized to protein input (Fig. 3B) or COX activity (Fig. 3C). It should be noted that, similar to RCR results, *S. subconica* and KEWJ had lower protein-normalized respiration rates in all respiratory states, although these differences were not statistically significant (Fig. 3B, min P = 0.13). Interestingly, KEWI had one of the highest protein-normalized respiration rates (Fig. 3B) but had the lowest COX-normalized rate in all states, but these differences were also not statistically significant (Fig. 3C, min P = 0.18).

Calculation of internally-controlled respiratory flux control factors (FCFs) (Table 2) revealed intra- and interspecific differences that were otherwise concealed by variability in mt quantity or quality among samples within a group. FCFs normalize flux to preceding reference states within each sample experiment, which enables evaluation of how specific ETS components contribute to respiratory control. The first FCF investigated was OXPHOS coupling efficiency, which is mathematically similar to the RCR and estimates the extent of ADP control over respiration. As with the RCR, KEWJ and *S. subconica* had OXPHOS coupling efficiencies that were lower than the other populations (Fig. 4A, *P* < 0.041). The proportion of NADH-supported OXPHOS contributed by external NADHs (DH_ex_) was higher in *S. subconica* compared to all **S. conica** accessions except KEWJ (Fig. 4B, *P* < 0.030). The CI FCF estimates the proportion of maximal OXPHOS contributed by CI and did not vary statistically among the groups (Fig. 4C, *P* > 0.652). However, the analogous CII FCF was higher in *S. subconica* than in **S. conica** ABR and KEWI (Fig. 4D, *P* < 0.020). The AOX FCF was not statistically variable among the accessions (Fig. 4E, *P* > 0.768), nor was excess CIV capacity (Fig. 4F, *P* > 0.193).

**Fig. 4.**
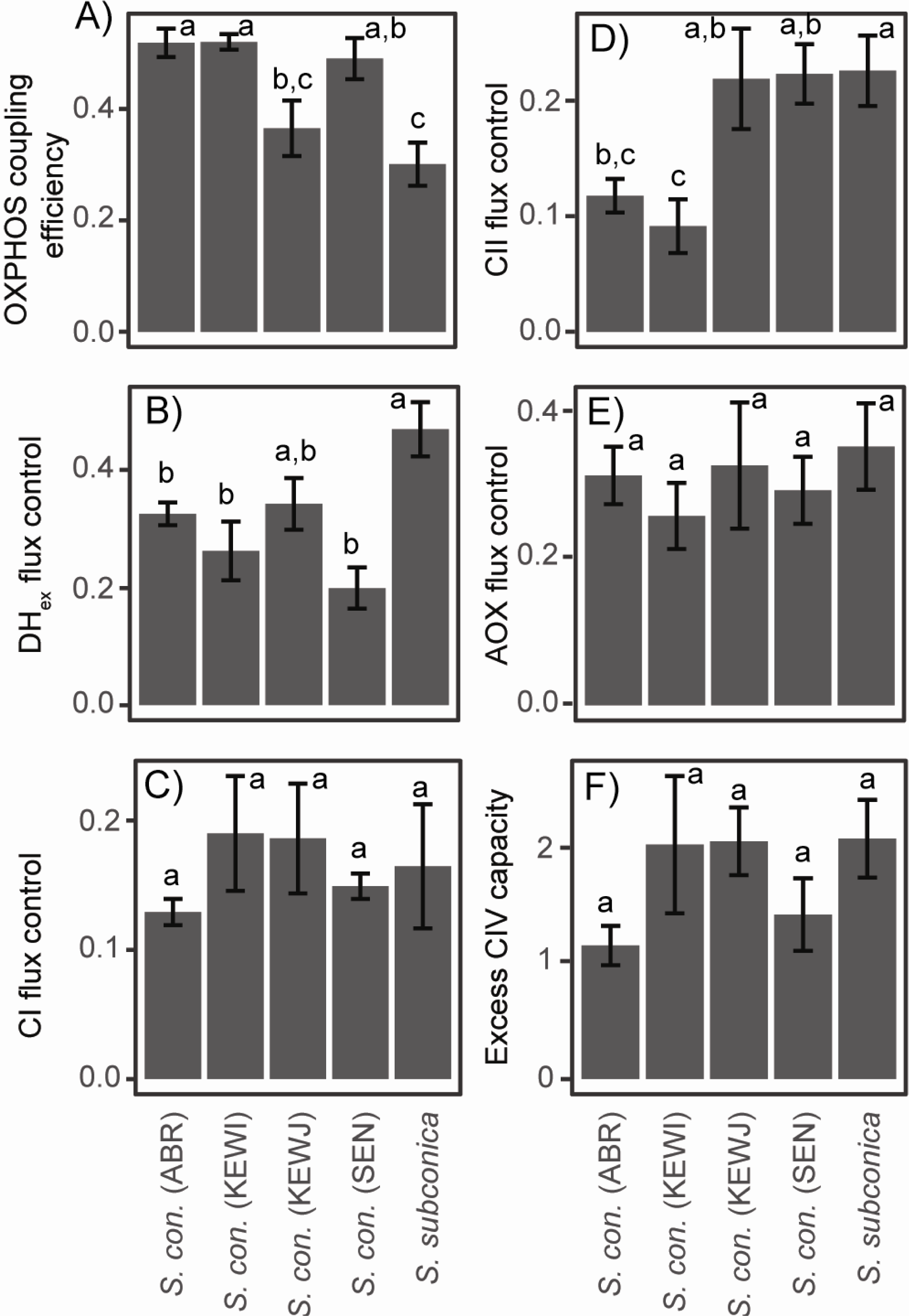
Variability in flux control factors (FCFs) among *Silene* accessions. See Table 2 for a description of FCFs. Lowercase letters indicate significant groupings among accessions at *P* < 0.05 based on Tukey post-hoc tests. Error bars show SEM. *n* = 7-15.

COX enzymatic activity when normalized to protein input was higher in **S. conica** KEWI and SEN compared to S.conica ABR and KEWJ (Fig. S5; 0.01 < *P* < 0.05). COX enzymatic activity was positively related to excess CIV capacity, although not statistically significant (P = 0.063, *r* = 0.26). COX activity was significantly and positively correlated with maximal OXPHOS flux (step D in Table 1 and Fig. 1: P = 0.048, *r* = 0.28), consistent with its central role in mt respiration.

### Correlations between FCFs and fitness metrics

Several FCFs were significantly correlated with each other (Fig. 5 and Table S2), with a few key themes emerging. First, OXPHOS coupling efficiency was negatively correlated with both DH_ex_ and CII flux, which were positively correlated with one another (Fig. 5, *P* < 0.016). Secondly, CI flux was positively correlated with CII and AOX flux (although not statistically significant: Fig. 5, P = 0.052 and 0.080, respectively), which were also positively correlated with one another (P = 0.002). Finally, in some cases these correlations are likely driven by differences between species. For example, *S. subconica* showed large CII and DH_ex_ fluxes, but low CI flux and OXPHOS coupling efficiency, while ABR showed the opposite pattern (Figs. 4 and 5).

**Fig. 5.**
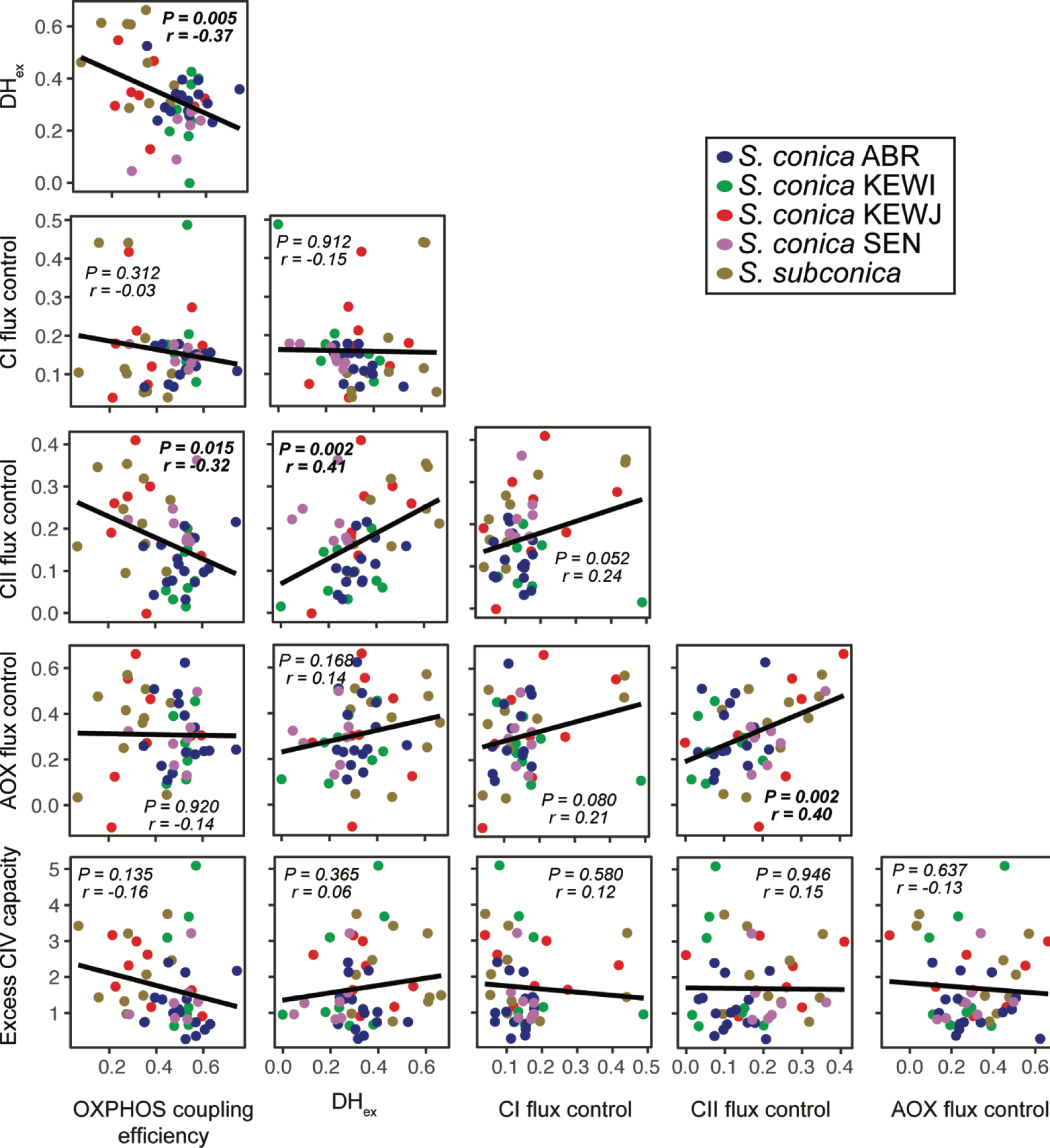
Correlation matrix between different flux control factors (see Table 2). P values are based on general linear models.

We also determined whether the plant fitness metrics quantified here were predictive of any FCFs or the maximal, protein-normalized respiration rate. Only a single correlation between an FCF and a plant fitness metric was statistically significant (Table S2) but was not so after correcting for multiple comparisons. There was a consistent positive correlation between all fitness metrics and maximal, protein-normalized respiration (Fig. S6; 0.03 > *P* > 0.90).

### Morphology and dynamics of *Silene* mitochondria

*Silene* seedling mitochondria are readily stained with TMRM allowing a rapid means to investigate their morphology and dynamics in living tissue (Fig. 6). The mitochondria of all three species investigated (*S. vulgaris, noctiflora*, and *latifolia*) show similar morphology (size and shape): physically discrete, spherical to short vermiform structures, typically 0.5 to 1 μm in diameter/length (Fig. 6, Fig. S7). As such, the mitochondria of these three species are morphologically similar to mitochondria in model species such as Arabidopsis and tobacco that have been visualised with TMRM, Green Fluorescent Protein, or the lipophilic fluorophore DiOC6 (Logan and Leaver, 2000; Van Gestel and Verbelen, 2002; Schwarzlander *et al*., 2012). Although a thorough quantification is beyond the scope of this work, other characteristics, such as number of mitochondria per cell, and their dynamic movement (Supplemental Videos S1 and S2) do not appear distinct from those of the model plant Arabidopsis.

**Fig. 6.**
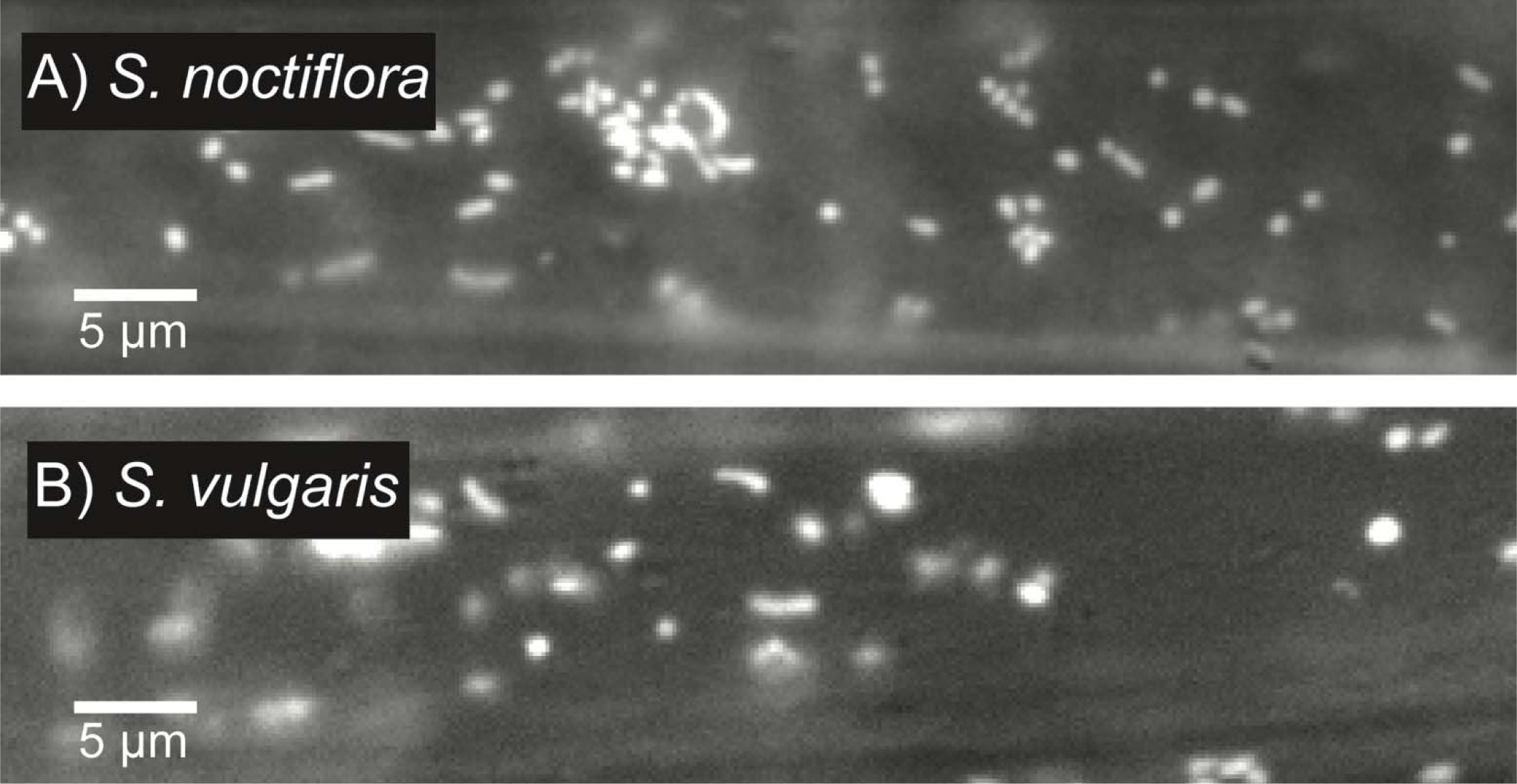
Epifluorescence micrographs of mitochondria in epidermal cells in intact roots of A) *Silene* noctiflora or B) *S. vulgaris* stained with TMRM. Each still image is taken from the respective Supplemental Videos S1 and S2. Bar = 5 µm.

## Discussion

### Do unusual mt genomes cause unusual mt function? Sometimes…

The discovery of multiple, independent angiosperm lineages that show elevated rates of sequence evolution and odd structures in their mt genomes (Mower *et al*., 2007) raises the question of whether mt function in these lineages has been altered. Here, we lay the groundwork for answering this question by examining two species in the genus *Silene* that display unusual mitogenomic features. Specifically, mt rates of evolution are highly elevated in these species (Sloan *et al*., 2009) − for example, there are 32 amino acid substitutions in Cox1 between Arabidopsis and **S. conica**, while there are only 7 substitutions between *A. thaliana* and *S. latifolia* (a close relative of **S. conica** with typical mt genomic features of angiosperms). These species also have incredibly expanded mt genomes (i.e., 11 Mbp vs. 0.3 Mbp) that have become fragmented into dozens of chromosomes (vs. a single chromosome)(Sloan *et al*., 2012a). Therefore, it is reasonable to imagine that either due to increased divergence at the molecular level or genomic rearrangements, mt function may be altered in these species. Here, we show that mt respiration in these species generally follows the expected patterns observed in other angiosperms with more typical mt genomes, and we develop a protocol to address the question in detail.

When substrates linked to CI, DH_ex_, and CII were added to isolated mitochondria, respiration increased as it has been shown to do in other angiosperms with typical mt genomes (Fig. 4; e.g, Douce *et al*., 1977). In addition, as in other angiosperms, inhibitors for CI and the AOX caused reduced respiration (Fig. 4; Ikuma and Bonner, 1967; Siedow and Girvin, 1980), while ASC/TMPD resulted in a large increase in respiration reflecting a significant excess capacity of cytochrome oxidase (Day *et al*., 1978). These results suggest CI, CII, CIV, DH_ex_, and AOX are functional in **S. conica** and *S. subconica* and likely play similar roles as in other angiosperms.

Importantly, our study did not include paired mt samples from closely related species such as *S. latifolia* and *S. vulgaris* that possess more “normal” plant mt genomes, which is an obvious next step for future studies. However, a first approach at visualizing mitochondria in *Silene* species with unusual patterns of mt genome evolution did suggest that abnormal mt genomic evolution does not lead to obviously abnormal mitochondria (Fig. 6). Further visualization experiments that pair SYBR Green and TMRM staining would be useful to determine whether mtDNA is distributed differently in the mitochondria of different *Silene* species. The range of RCRs observed here (1.1 −3.9) is also similar (1.3 − 3.7) to that reported for leaves from adults in peas (Flowers, 1974; Day *et al*., 1985). The response to substrates such as NADH and succinate were also comparable to those observed in other systems (Karapanos *et al*., 2009). Moreover, preliminary experiments performed while developing the mt respiration protocol using *S. latifolia* and *P. sativum* resulted in qualitatively similar results.

Calculating FCFs as performed here is also an attractive methodology for directly comparing species with unusual vs. typical mt genomes. FCFs allow specific respiratory components to be quantified by normalization to a common reference respiratory state. However, they have not been widely adopted in plant mt respiration experiments. By comparing FCFs instead of respiration rates, the amount (and to a lesser extent, quality) of mitochondria loaded in a particular sample is controlled for, making this approach ideal for comparing crude mt isolates from even distantly related species. Specifically, when comparing species with normal vs. atypical mt genomes, we might predict that FCFs that capture OXPHOS-linked respiration contributed by chimeric mitonuclear OXPHOS complexes (e.g., CI and CIV) would vary, while FCFs that describe flux mediated by complexes composed solely of nuclear-encoded subunits (e.g., DH_ex_, CII, and AOX) would serve as controls. Therefore, while it is not possible at this time to provide a detailed quantification of mt respiration in *Silene* species with unusual vs. typical mt genomes, our current data do not indicate any major alterations in mt form or function in *Silene* species with unusual patterns of mt evolution.

Why do some lineages with fast-evolving mt genomes such as *Viscum* have altered mt function, while *Silene* appears to have largely typical patterns of mt respiration? Accelerated evolutionary rates in *Viscum* are associated with the loss of many mt genes and a large reduction in genome size (Skippington *et al*., 2015, 2017). Importantly, CI mt genes have been entirely lost in *Viscum* (i.e., not transferred to the nuclear genome), which explains why CI activity has been lost and activities of the AOX and alternative NADH dehydrogenases are elevated (Maclean *et al*., 2018). In addition, the formation of OXPHOS “supercomplexes” and mt morphology are altered in Viscum, also likely due to loss of CI (Senkler *et al*., 2018). On the other hand, accelerated mt evolution in *Silene* has resulted in expanded genomes and retention of protein coding genes. Moreover, signals of purifying selection (i.e., d_N_/d_S_ << 1) from mt-encoded OXPHOS genes in *Silene* with accelerated mt rates were as strong as those in species with slow rates of mt evolution (Havird *et al*., 2015). Therefore, while both *Viscum* and *Silene* have experienced patterns of accelerated evolution in their mt genomes, the types of evolution (relaxed vs. purifying) and implications for mt function are not the same. We speculate that this may be due to the parasitic vs. free-living lifestyles of *Viscum* v*S. *Silene**. Although not all parasitic plants have reduced mt gene content (Fan *et al*., 2016), they do tend to show elevated rates of mt genomic evolution compared to free-living relatives (Bromham *et al*., 2013), possibly reflecting increased mutation rates or relaxed selection.

### Inter-and intraspecific variation in *Silene* mt function

There was significant inter-and intraspecific variation in the mt properties examined here. Most notably, *S. subconica* and *S. conica* KEWJ had lower OXPHOS coupling efficiency (Fig. 4A), lower overall respiration rates (Fig. 3B), greater CII control over respiration (Fig. 4D), and lower COX activity (Fig. S5) than other accessions of **S. conica**. One explanation for these results is that specific populations or species of *Silene* may have undergone environmental adaptation in mt function. For example, in Plantago, another angiosperm genus that shows variation in rates of mt genome evolution, closely related highland and lowland species had variable rates of leaf respiration (Atkin *et al*., 2006). Respiration also varies in general across climates and geography for a wide range of species (Atkin *et al*., 2015). Both *S. conica* and *S. subconica* have large distributions with somewhat isolated populations found in disparate environments − from mountain tops to sandy beaches (Jalas and Suominen, 1988). Therefore, it is possible that specific populations have adapted to utilize slightly different mt respiratory pathways. Given the high rate of molecular evolution in the mt genomes of these species, the phenotypic variation observed here might even be partly attributable to variation in mtDNA.

Another reason for the inter-and intraspecific variation observed here may be that, despite our best efforts to control environmental factors in the greenhouse, some accessions may have experienced slightly different environments. It has long been known that mt function is plastic in plants. For example, pea seedlings grown under water stress had lower RCRs than controls (Generozova *et al*., 2009) and AOX flux control also varies with environment (Fiorani *et al*., 2005; Vanlerberghe, 2013; Ahanger *et al*., 2017). Given our relatively low sample sizes and the large amount of within-accession (i.e., within genotype) variation seen in basic plant fitness metrics (Fig. 2), it is possible that differences in watering and microenvironmental conditions may explain some of the variation observed here. Future experiments performed in climate-controlled growth chambers may alleviate this source of variation.

### A novel protocol for assessing mt respiration

One of our primary goals was to develop a novel respirometry protocol that investigates the relative contributions of specific ETS components to plant mt respiration. The protocol developed here (Fig. 1A) does not use any novel substrates or inhibitors, but the order in which they are added to the reaction allows for a detailed accounting of FCFs capable of elucidating inter-and intraspecific variation in mt function not revealed by conventional assays. Cross-species analyses in the present study found that mitochondria with low OXPHOS coupling efficiency tended to have higher CII, DH_ex_, and AOX fluxes, suggesting a greater reliance on these latter components over CI (Fig. 5). These inverse relationships make sense, because CII, DH_ex_, and AOX do not couple electron transfer with proton translocation that powers the ATP synthase, whereas CI does. This type of “inefficient” respiratory profile has been observed previously in *Nicotiana sylvestris* showing cytoplasmic male sterility due to mitonuclear incompatibility (see below; Sabar *et al*., 2000). Samples from *S. subconica* and to a lesser extent the **S. conica** KEWJ exhibited this profile, while the other samples showed greater respiratory control by CI. Overall respiration, which was lower in *S. subconica*, was also correlated with fitness (Fig. S6), suggesting that different respiratory profiles in *Silene* may be under selection if they have a genetic basis.

A promising application of our protocol concerns the concept of mitonuclear incompatibility (Hill, 2017; Sloan *et al*., 2017). Our results confirm that complexes composed entirely of nuclear-ߝencoded subunits as well as chimeric complexes with mt-encoded subunits as their core (Fig. 1A) both contribute to mt respiration in these *Silene* species. Under a mitonuclear (in)compatibility framework, mt and nuclear genomes within a population or species are thought to be “matched” to one another due to a long history of coevolution. However, when species or populations interbreed, resulting offspring can end up with “mismatched” genomes from different lineages, resulting in reduced fitness in hybrids and ultimately reproductive isolation between populations (Burton and Barreto, 2012; Hill, 2016; Sloan *et al*., 2017). Chimeric mitonuclear OXPHOS complexes containing mt-encoded subunits are hypothesized to be affected by compromised mitonuclear interactions, while complexes composed entirely of nuclear-encoded subunits should not be affected. Our protocol and the FCFs calculated here allow mt respiration to be parsed into contributions from chimeric and purely nuclear complexes and should therefore allow this hypothesis to be tested in a novel way using systems showing compromised mitonuclear interactions (e.g., Burton *et al*., 2006).

Angiosperms and *Silene* in particular offer unique opportunities to study the functional implications of mitonuclear incompatibilities. In animal-based studies, CII is the only OXPHOS complex composed solely of nuclear-encoded subunits and has therefore become the “go-to” control for mitonuclear incompatibilities in studies of mitonuclear coevolution and mt function (Ellison and Burton, 2006). Plants offer an additional set of nuclear-encoded controls in the AOX and alternative NADH dehydrogenases. Additionally, CII is entirely nuclear-encoded in some angiosperms but has retained some mt-encoded subunits in other lineages (Adams *et al*., 2002). Therefore, mitonuclear incompatibilities induced by hybridization should produce different but predictable effects on mt function between plants and animals and among different angiosperm lineages. Results presented here and elsewhere (Sabar *et al*., 2000) suggest that the proportion of respiration contributed by the AOX, alternative NADHs, and CII may be elevated in systems with compromised mitonuclear interactions, perhaps reflecting a mismatch between substrate oxidation and ADP phosphorylation capacity. *Silene* in particular offers an interesting system to begin to examine these effects, as clear evidence for mitonuclear coevolution has been found using different *Silene* lineages with variable rates of mt evolution (Sloan *et al*., 2014; Havird *et al*., 2015; Havird *et al*., 2017). A clear future goal for studies of mitonuclear interactions would therefore be to examine hybrids predicted to show mitonuclear incompatibilities with a protocol like the one developed here.

## Acknowledgements

We wish to thank Lance Li Puma for assistance with the O2K, members of the Graham Peers lab for assistance with COX activity assays, and members of the Sloan lab for comments on this work. This work was supported by NIH F32GM116361 to JCH and NSF MCB 1412260 to DBS.

## Supplementary data

Table S1. Details on substrates, cofactors, and inhibitors used in the mt respiration protocol.

Table S2. Correlations among metrics of plant fitness and flux control factors (Table 2). P values are presented above the diagonal (based on linear models), while r2 values are presented below the diagonal. Significant correlations at *P* < 0.05 are bolded.

Fig. S1. Mitochondrial respiration in *Silene* conica (SEN) during the protocol detailed in Fig. 1, showing that peak values following the addition of ADP (D) are generally recovered if given enough time.

Fig. S2. Correaltion between total seed mass and the number of capsules.

Fig. S3. Mitochondrial respiration under increasing malate concentration (M) in *S. noctiflora* and *P. sativum*.

Fig. S4. Respiratory control ratios (RCR) in *Silene*.

Fig. S5. Cytochrome c oxidase (COX) activity in *Silene* normalized to protein input.

Fig. S6. Correlations between plant fitness metrics and maximal, protein normalized respiration rate.

Fig. S7. Epifluorescence micrographs of mitochondria in an epidermal cell of an intact root of *Silene* latifolia stained with TMRM.

Supplemental Video S1. Movie of mitochondria in an epidermal cell in an intact root of *Silene* noctiflora stained with TMRM.

Supplemental Video S2. Movie of mitochondria in an epidermal cell in an intact root of *Silene* vulgaris stained with TMRM.

## Abbreviations

ABR/SEN/KEWI/KEWJ, ***S. conica*** accessions used here; AOX, alternative oxidase; ASC, ascorbate; BSA, bovine serum albumin, CI-CV, Complexes I-V; COX, cytochrome c oxidase; CytC, cytochrome c; DH_in_ and DH_ex_, internal and external NADH dehydrogenases; ETS, electron transfer system; FCFs, flux control factors; mt, mitochondrial; OXPHOS, oxidative phosphorylation; nPG, npropyl gallate; O2K, Oxygraph 2K; RCR, respiratory control ratio; ROT, rotenone; SUCC, succinate; TMPD, tetramethylphenylenediamine; TMRM, tetramethylrhodamine methyl ester

## Abbreviations

ABR/SEN/KEWI/KEWJ: S. conica accessions used here
AOX: alternative oxidase
ASC: ascorbate
BSA: bovine serum albumin
CI-CV: Complexes I-V
COX: cytochrome c oxidase
CytC: cytochrome c
DH_in_ and DH_ex_: internal and external NADH dehydrogenases
ETS: electron transfer system
FCFs: flux control factors
mt: mitochondrial
OXPHOS: oxidative phosphorylation
nPG: n-propyl gallate
O2K: Oxygraph 2K
RCR: respiratory control ratio
ROT: rotenone
SUCC: succinate
TMPD: tetramethylphenylenediamine
TMRM: tetramethylrhodamine methyl ester

